# Fluvial seeding of cyanobacterial blooms in oligotrophic Lake Superior

**DOI:** 10.1101/2020.11.03.366955

**Authors:** Kaitlin L. Reinl, Robert W. Sterner, Brenda Moraska Lafrancois, Sandra Brovold

**Affiliations:** Large Lakes Observatory, University of Minnesota Duluth, Duluth, MN 55812, USA; National Park Service, Ashland, WI, USA

**Keywords:** Cyanobacteria, algal loading, oligotrophic, harmful algal blooms

## Abstract

Lake Superior has recently begun experiencing cyanobacterial blooms comprised of *Dolichospermum lemmermannii* near the Apostle Islands and along the southern shore of the western arm. Little is known about the origin of these blooms. Experiments were conducted during the summers of 2017 and 2018 to identify sources of propagules and characteristics of sites that were potential sources. The 2017 experiments were conducting using a factorial design with three source zones (Harbor, River, and Lake), two nutrient conditions (high and low N:P), and three temperatures (15, 20, and 25 °C). At the end of the experiment, cyanobacteria were most abundant from the ‘River’ and ‘Harbor’ zones at low N:P and 20 and 25 °C, with *D. lemmermannii* most abundant at 20 °C. Subsequently in 2018 we evaluated 26 specific inland locations from three waterbody types (Coastal, Lake/Pond, and River) and explored similarities among those sites that produced cyanobacteria in high abundance when samples were incubated under optimal conditions (low N:P and 25 °C). Under these growing conditions, we found high cyanobacteria abundance developed in samples from river sites with low ambient temperatures and high conductivity. Field monitoring showed that Lake Superior nearshore temperatures were higher than rivers. These observations suggest that blooms of *D. lemmermannii* in Lake Superior are initiated by fluvial seeding of propagules and highlight the importance of warmer temperatures and favorable nutrient and light conditions for subsequent extensive cyanobacterial growth. We argue that the watershed is an important source of biological loading of *D. lemmermannii* to Lake Superior, and that when those cells reach the nearshore where there is warmer water temperatures and increased light, they can grow in abundance to produce blooms.

## 1. Introduction

Cyanobacterial blooms are a major threat to the beneficial use of marine and freshwater ecosystems globally. Along with the rise in the spatial and temporal extent of nuisance cyanobacterial blooms (Huisman et al., 2018; Taranu et al., 2015), there has also been a rise in the number of blooms that are toxic (Anderson, 1989; Sukenik et al., 2015), posing a significant threat to public health. It is widely recognized that cyanobacterial blooms occur in four of the five Laurentian Great Lakes (LGL); however, less well known is that localized surface blooms of *Dolichospermum lemmermannii* (formerly known as *Anabaena lemmermannii* (Wacklin et al., 2009) have been observed along the southern shore of the western arm in oligotrophic Lake Superior. In 2012 and 2016-2018, cyanobacterial blooms were detected in the nearshore with varying size and duration, most of which were very localized and dissipated quickly; however, in 2018 the bloom persisted for approximately one week in early August and extended approximately 100 km alongshore from Duluth to the Apostle Islands region and 3 km offshore with the highest densities being observed at the shoreline (Sterner et al., submitted for review).

Blooms of cyanobacteria are often associated with increased loading of nutrients, particularly phosphorus (Lürling et al., 2018; Schindler, 1975; Steinberg and Hartman, 1988); however, Lake Superior is a cold oligotrophic system that is low in phosphorus (Sterner, 2010). The watershed in the region where algal blooms have been observed in Lake Superior is primarily forested, but even small differences in land cover and position influence nearshore water quality (Yurista et al., 2011). The western arm has become increasingly subject to extreme precipitation events delivering large amounts of sediment and nutrients to the nearshore (Cooney et al., 2018). Two 500-1000 year rainfall events that occurred on the south shore of Lake Superior in 2012 and 2018 (Cooney et al., 2018; Roache et al., 2020) were followed by cyanobacterial blooms varying in size and duration with a lag of 25 days in 2012 and 53 days in 2018; however, small blooms have also occurred in 2016 and 2017 without such major rain events. Extreme storms provide large fluxes of nutrients and organic material to the lake (Cooney et al., 2018; Kling et al., 2000; Minor et al., 2014). Cyanobacterial blooms are also generally associated with higher water temperatures (Paerl and Huisman, 2008), and Lake Superior has been warming at the fastest rate of the LGL by maximum summer surface temperature (Austin and Colman, 2007), which may be contributing to the emergence of cyanobacterial blooms (Sterner et al., submitted for review). As summer epilimnetic temperatures increase, conditions can favor cyanobacteria and result in subsequent blooms (Konopka and Brock, 1978; Kosten et al., 2012; Paerl and Huisman, 2009; Robarts and Zohary, 1987).

Coastal regions of both marine and freshwater systems can be hotbeds for cyanobacterial blooms (Cloern, 1996; Cloern and Jassby, 2010, 2008). Coastal regions are transitional areas that are significantly influenced by watershed inputs as well as exchange with offshore waters (Cloern, 1996; Howell et al., 2012). Thus, consideration of the surrounding landscape is critical to understanding bloom development in coastal areas (Kratz et al. 1997).

Rivers are often recognized as sources of nutrients for algal growth in lakes, but seldom are inflowing waters also considered as sources of the living cells that initiate blooms. The riverine input of propagules to receiving lakes is referred to as “fluvial seeding” and the importance of such seeding is referred to as the algal loading hypothesis (ALH) (Conroy, 2007; Loftin et al., 2016). Discussion of the ALH so far is a relatively unexplored question (Conroy et al., 2008), and nearly all of the work done regarding this hypothesis has been in Lake Erie and its watershed, despite any evidence that this process is be unique to that system (Bridgeman et al., 2012; Conroy et al., 2014; Davis et al., 2014; Kutovaya et al., 2012). Conroy et al. (2014) conducted a study evaluating major tributary inputs as sources of seed populations of cyanobacteria propagules to the Lake Erie blooms and found elevated concentrations of *Microcystis* in the rivers and observed visible riverine blooms, concluding that the rivers were playing a key role in the delivery of seed populations and bloom material to Lake Erie. In contrast, studies using DNA fingerprinting techniques have found that the cyanobacterial bloom communities found in the lake are not very similar to those sampled in the tributaries (Chaffin et al., 2014; Kutovaya et al., 2012). The disagreement in findings on the role of fluvial seeding in Lake Erie demonstrates the lack of knowledge regarding these processes and illustrates the need for further research in this area. Answers to these questions are critical in effectively addressing current and future threats of cyanobacterial blooms in coastal waters.

Despite being episodic and relatively small in spatial scale, algal blooms in Lake Superior have generated public concern due to their unprecedented nature and their potential impacts on public health, aesthetics, and the local tourism economy. In this work we explore how land-lake interactions relate to the emergence of cyanobacterial blooms in Lake Superior and provide insights to mechanisms responsible for their appearance. Specifically, we investigate the question - What are the sources of propagules leading to cyanobacterial blooms in Lake Superior and are there similarities among locations that are potential sources?

## 2. Material and methods

Laboratory experiments were conducted in summer of 2017 and 2018 to identify potential sources of cyanobacterial blooms in western Lake Superior and to identify common characteristics of potential upstream sources. The 2017 experiment aimed to identify the major habitat types that are sources of cyanobacteria and determine whether Lake Superior blooms likely originate in the lake or were initiated in waterbodies further inland (Figure *1*). The 2018 experiment was informed by results from the 2017 experiments and examined 26 specific inland locations as potential sources of blooms. The 2018 experiments also identified some characteristics of sites with higher potential for cyanobacterial growth and thus likelihood of acting as a source for blooms in the large lake (Figure 1).

**Figure 1.**
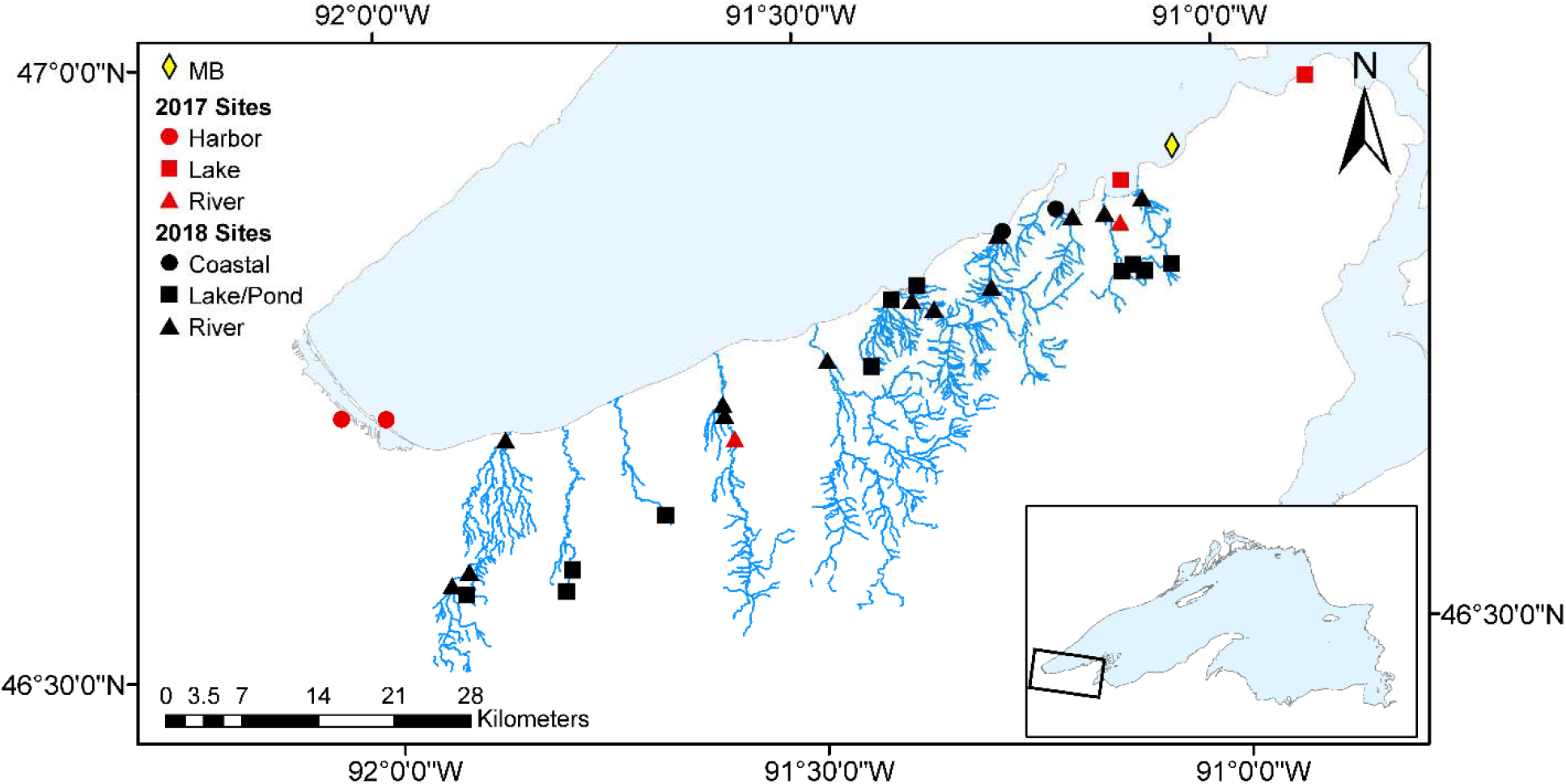
Map of study sites. Sites from 2017 are shown in red with shapes denoting the zone, and 2018 sites are shown in black with shapes denoting the waterbody type. Stream systems associated with 2018 sites are also shown. A yellow diamond denotes the location of the Mawikwe Bay (MB) field sampling site.

Experiments were designed to assess the propensity of a biological community from a given location to develop into a cyanobacterial bloom over ecologically realistic time scales. Thus, inocula were brought into the lab and incubated in a wide range of nutrient and temperature conditions capable of supporting cyanobacteria. The abundance of major algal groups was recorded over time. This experimental approach was chosen because it ensured that if a viable cyanobacterial population was present, we would observe growth during the experimental time frame. We concluded that inocula that did not develop blooms under bloom-promoting conditions within the experimental period were unlikely to be the source of bloom propagules *in situ.*

### 2.1 Sample Collection and pre-processing

We defined three “zones” or habitat types during the 2017 experiments: 1) Lake Superior near the Apostle Islands (‘Lake’), 2) inflowing rivers (‘River’), and 3) the Duluth-Superior harbor (‘Harbor’). Each of these three zones was comprised of two nearby locations: ‘Lake’ included two nearshore (max depth = 5-10m) locations in Lake Superior, ‘River’ included the Bois Brule and Siskiwit Rivers, and ‘Harbor’ included Wisconsin Point and Superior Bay (Figure 1). Water samples that were collected in each zone during the 2017 experiments were composited to increase chances of collecting viable cyanobacterial propagules. Chemical analyses were conducted on the composited water samples. Habitat types were defined differently in 2018. The 26 sites studied in 2018 included two coastal waters (characterized by influence from both inland rivers/streams and Lake Superior) (‘Coastal’), 11 upland lakes/ponds (‘Lake/Pond’), and 13 rivers (‘River’), and were selected for their accessibility and connectivity to Lake Superior.

Water, net tows, and sediment were collected from each location to be used as inoculant in the laboratory experiments. Whole surface water was collected using a clean, acid washed carboy and filtered through a 150 μm mesh to remove large debris. Horizontal surface net tows were collected with an 80-μm mesh plankton net, and surface sediment was collected via grab sample using a 50 ml centrifuge tube or first collected using a PONAR at deeper sites and then transferred to the centrifuge tube. Net tows were used to provide an additional concentrated inoculant.

Supernatant from sediment samples that had been allowed to settle for 1 minute after agitation was added to capture vegetative cells or akinetes that were in the sediment. Filtered water samples, net tows, and supernatant pipetted from sediment samples from each zone were composited with their respective sample and used as inoculant. The amount of each sample type in the inoculant was 500 ml water, 17 ml net tows, and 2 ml sediment supernatant in 2017, and 200 ml of water, 30 ml of net tow, and 10 ml of sediment supernatant in 2018. Temperature and conductivity were also measured from all sampling sites in July and August in 2018. Electrical conductivity (EC) was measured using an UBANTE™ total dissolved solids (TDS) meter in parts per million (ppm). The TDS data was converted to EC in mS/cm and then normalized to 25°C (EC_25_) to account for temperature effects using Eq (1) (Atkins and de Paula, 2006), where T is temperature in °C.

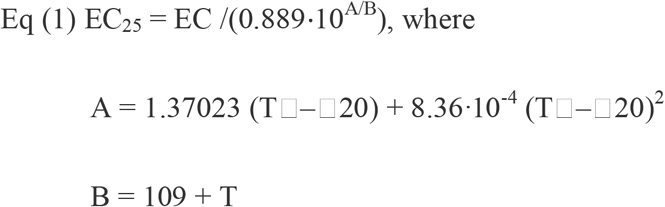

### 2.2 Chemical Analyses

Water chemistry for 2017 was analyzed for composited water samples from each zone (3 zones) and 2018 samples were analyzed for individual sites (26 sites). Particulate fractions of carbon (POC) and nitrogen (PON) were collected on 25mm pre-ashed GF/F filters and particulate phosphorus (PP) was collected on 25mm pre-ashed, acid-rinsed GF/F filters. POC and PON filters were then dried at 60°C until analysis on a Costech Elemental Analyzer. Chlorophyll-a (chl-a) samples were filtered using 25mm 0.2 μm cellulose nitrate filters, and frozen in the dark until analyzed according to Welschmeyer (1994) on a Turner Design 10-AU fluorometer after extraction in 90% acetone for 20-22 hours in the dark at 4 °C. Total dissolved phosphorus (TDP), soluble reactive phosphorus (SRP), dissolved organic carbon (DOC), ammonia (NH_3_), and soluble reactive silica (Si) samples were filtered through 0.22-μm filters, and frozen until analyzed, with the exception of DOC which was acidified to pH of 2. Total phosphorus (TP) and nitrate (NO_3_^−^), were collected as whole-water samples and then frozen for preservation. PP, TP, and TDP samples were digested using potassium persulfate (modification of EPA method 365.1) and then analyzed on a SEAL AQ400. SRP samples were analyzed on a SEAL AQ400 on undigested sample (EPA method 365.1, and Murphy and Riley (1962)). NO_3_^−^ and Si were analyzed according to EPA method 353.2 and Standard Methods 4500-SiO_2_, respectively, and also processed on a SEAL AQ400. DOC and TDN were analyzed using a Shimadzu TOC-Vcsh and a TNM-1 module. NH_3_ was analyzed according to (Taylor et al., 2007), modified from (Holmes et al., 1999), with the exception of samples that exceeded the upper limit for those methods, in which case they were analyzed on the SEAL AQ400 using EPA method 350.1. Total nitrogen (TN) and was calculated as the sum of TDN and PON, and total organic carbon (TOC) was calculated as the sum of DOC and POC.

### 2.3 Laboratory Conditions

In 2017, cultures were exposed to high and low nitrogen:phosphorus (TN:TP) nutrient conditions (high: 50:1, low:1.5:1) (all ratios are molar), and three temperatures (15, 20, and 25°C). This broad set of conditions was chosen to insure we would observe cyanobacterial growth, as well as to take a coarse look at ideal conditions, because the species of cyanobacteria observed in Lake Superior blooms has been observed in both cold, oligotrophic and warm, eutrophic waters. Cultures were run in triplicate for a total of 54 samples (3 locations x 2 N:P x 3 temperatures x 3 replicates). Cultures were incubated in 1-L polycarbonate bottles (previously acid washed), continuously bubbled slowly with room air, and exposed to approximately 250 μmol photons/m^2^/s for 24 hours per day for 21 days. However, the 15°C incubator failed 3 days prior to the scheduled end of the experiment, and the last growth measurement was taken 3 days prior to that; therefore, no data were used for the 15°C treatment in the final 6 days of the experiment, including samples for taxonomic identification. The experiment began with 100% environmental sample (500 ml water, 17 ml net tows, and 2 ml sediment supernatant) and then was run as a semi-continuous culture, with 150 ml of the sample volume being exchanged for new sterile media every other day, after thoroughly swirling the flasks. Culture flasks were exchanged to discourage wall growth when visible. Cultures were preserved using Lugol’s fixing solution (Wetzel and Likens, 2000) and were examined for phytoplankton identification and counts (cells and colony units) by the Wisconsin State Lab of Health and Hygiene, Madison, WI. Filtered and acetone-extracted chl-a concentration was also measured in each culture at the end of the experiment.

In 2018, cultures were incubated under the same conditions as in 2017 with the following modifications. In 2018 only the low N:P and 25°C conditions were used, as they proved to be the best conditions for cyanobacterial growth in the previous year’s experiment. Cultures were also run as batches, duplicates were used instead of triplicates due to space limitations, and the experiment was run for 10 days. Each bottle received 700 ml of low N:P medium and was inoculated with 200 ml of water, 30 ml of net tow, and 10 ml of sediment supernatant. In 2018, a subset of 14 samples from sites with the highest cyanobacteria biomass, as measured by the PHYTO-PAM (Walz, 2003), were examined for identification and counts due to cost limitations.

### 2.4 Growth Media

The medium used in these experiments was originally designed for an experiment aimed at evaluating competition in natural algal communities in western Lake Superior, western Lake Erie, and Lake Huron, but here we only report its use in experimental work for Lake Superior. The Average Laurentian Great Lakes (A-LGL) Medium was developed for this work based on WC (Guillard, 1975), COMBO (Kilham et al., 1998), SuFr (Twiss et al., 2004), and HH-COMBO (Baer and Goulden, 1998) media, as well as major ion concentrations in the Laurentian Great Lakes (LGL) (Chapra et al., 2012). Table 1 shows the total concentration of major ions in several media and the LGLs. Each of the media were compared to LGL major ion concentrations and used to create a new media recipe, optimized for meeting LGL averages.

**Table 1.** Major ion concentrations of selected algal growth media and LGLs. A-LGL was used in this study. All concentration reported in μmol/L. Ca^2+^ estimates include CaCO_3_.

| **Major Ions** | **$\text{Ca}^{2+}$** | **$\text{Mg}^{2+}$** | **$\text{Na}^+$** | **$\text{K}^+$** | **$\text{SO}_4^{2-}$** | **$\text{Cl}^-$** | **$\text{HCO}_3^-$** | **Total Ions** |
| --- | --- | --- | --- | --- | --- | --- | --- | --- |
| WC | 250 | 150 | 1373 | 100 | 150 | 5367 | 150 | 7540 |
| COMBO | 250 | 522 | 150 | 1374 | 200 | 150 | 150 | 2796 |
| SuFr | 335 | 100 | 88 | 26 | 60 | 480 | 13 | 1102 |
| HH COMBO | 750 | 225 | 1513 | 100 | 225 | 1522 | 1500 | 5835 |
| A-LGL | 600 | 200 | 214 | 25 | 200 | 1238 | 150 | 2627 |
| Superior | 758 | 116 | 63 | 13 | 40 | 40 | 834 | 1864 |
| Michigan | 1974 | 464 | 272 | 36 | 250 | 340 | 2140 | 5476 |
| Huron | 1442 | 307 | 168 | 24 | 165 | 186 | 1563 | 3855 |
| East Erie | 1688 | 366 | 373 | 37 | 237 | 411 | 1758 | 4870 |
| Ontario | 1736 | 354 | 503 | 38 | 266 | 552 | 1789 | 5238 |
| LGL Average | 1520 | 321 | 276 | 30 | 192 | 306 | 834 | 3479 |

**Table 2.** A-LGL medium recipe.

| Major Stocks | $\mu\text{mol/L}$ |
| --- | --- |
| $\text{CaCl}_2 \cdot 2\text{H}_2\text{O}$ | 600 |
| $\text{MgSO}_4 \cdot 7\text{H}_2\text{O}$ | 200 |
| $\text{NaHCO}_3$ | 150 |
| $\text{Na}_2\text{SiO}_3 \cdot 5\text{H}_2\text{O}$ | 20 |
| $\text{H}_3\text{BO}_3$ | 388 |
| KCl | 25 |
| Algal Trace | $\mu\text{mol/L}$ |
| $\text{Na}_2\text{EDTA}$ | 14.92 |
| $\text{FeCl}_3 \cdot 6\text{H}_2\text{O}$ | 3.70 |
| $\text{CuSO}_4 \cdot 5\text{H}_2\text{O}$ | 0.01 |
| $\text{ZnSO}_4 \cdot 7\text{H}_2\text{O}$ | 0.08 |
| $\text{CoCl}_2 \cdot 6\text{H}_2\text{O}$ | 0.08 |
| $\text{MnCl}_2 \cdot 4\text{H}_2\text{O}$ | 0.91 |
| $\text{Na}_2\text{MoO}_4 \cdot 2\text{H}_2\text{O}$ | 0.09 |
| $\text{H}_2\text{SeO}_3$ | 0.01 |
| $\text{Na}_3\text{VO}_4$ | 0.01 |
| Vitamins | $\mu\text{mol/L}$ |
| Thiamine · HCl | 296.5 |
| Biotin | 0.20 |
| $\text{B}_{12}$ | 0.037 |
| Nutrients | $\mu\text{mol/L}$ |
| $\text{K}_2\text{PO}_4$ (High N:P) | 0.5 |
| $\text{K}_2\text{PO}_4$ (Low N:P) | 5.33 |
| $\text{NH}_4\text{NO}_3$ (High N:P) | 25 |
| $\text{NH}_4\text{NO}_3$ (Low N:P) | 8 |

### 2.5 Growth Measurements

Due the large number of samples in the growth measurements, a PHYTO-PAM (Walz, GmbH) was used to monitor growth. The PHYTO-PAM was used with the Phyto-Win software (V 1.45) and the emission detector (ED) unit. The PHYTO-PAM employs multi-spectral fluorescence techniques to differentiate between “blue-green”, “green”, and “brown” algal groups which generally correspond to cyanobacteria, chlorophytes, and diatoms, respectively. High frequency pulses of light are emitted and the excitation of chlorophylls at different wavelengths are measured to separate algal groups. This measurement has proven to provide reliable information on abundance of major groups, provided certain precautions are taken and samples are validated (Lürling et al., 2018; Walz, 2003). To measure growth in our samples, each bottle was subsampled and left in the dark for approximately 30 minutes to be dark-adapted, and all PHYTO-PAM measurements were taken in the dark. Each sample was pipetted into a clean cuvette and placed in the ED unit. The gain was adjusted for each sample, and the amount of chlorophyll in each algal group was measured using chlorophyll measuring frequency (Chl (MF)) mode with a measuring frequency of 32.

The PHYTO-PAM, like virtually all measures of chlorophyll, is imperfect. Measurements rely on proper reference values being supplied to the software for the samples being analyzed, which can prove complicated when dealing with natural samples with an unknown community composition. To combat this uncertainty, we took the following steps. First, we validated measurements on the PHYTO-PAM using pure cultures of *Microcystis*, *Ankistrodesmus*, and *Cyclotella*, to confirm that the PHYTO-PAM was correctly identifying “blue-green”, “green”, and “brown” algal groups, respectively (Figure S1). We also preserved samples at the end of each experiment for identification. Finally, we treated PHYTO-PAM measurements as a relative measure, meaning that we did not interpret absolute values, but instead evaluated relative changes over time (growth). Though the PHYTO-PAM should not be relied on to give precise biomass estimates, it can provide useful information on the abundances of major algal groups (Lürling et al., 2018).

### 2.6 Data Analysis

A 3-way MANOVA was used to test for differences in the response of cyanobacterial growth in the 2017 experiments. Each treatment was a different factor (temperature, zone, and N:P) and the response variables were growth rates of the blue-green, green, and brown algal groups, but because the last 15°C treatment measurements were taken 6 days prior to the end of the experiment, only the measurements up to day 13 were used to calculate growth rates for the 15°C treatment. Growth rates were calculated as the slope of the natural log of PHYTO-PAM measurements over time. Growth rates were calculated for the full duration of the experiment for all treatments as well as up to day 13 to determine if there were differences in results under the two windows of time and none were found. We also tested for differences among treatments using taxonomic observations of the relative abundance of *D. lemmermannii* by applying the non-parametric Kruskal-Wallis and Kruskal-Wallis multiple comparison (adjusted to avoid type I errors (Siegel and Castellan, 1988) tests using the *stats* and *pgirmess* packages in R (CoreTeam, 2017), respectively.

Due to the spatial nature of the 2018 data, we tested for spatial autocorrelation in site characteristics and experimentally derived growth rates by calculating Moran’s I (Moran, 1950) using the inverse distance squared method in ArcMAP (V 10.4.1). Moran’s I tests for tendency of units to be similar to its neighbors by calculating a correlation coefficient ranging from −1 to 1, where values near 1, 0, and −1, are clustered, random, or dispersed, respectively. We found that the response variable was not spatially autocorrelated, but many of the potential predictors were autocorrelated as clusters. We then conducted a visual assessment of semivariograms with varied correlation structures to assess severity of autocorrelation, and concluded that corrective measures were not required. We applied paired two-sided t-tests to test for differences in site characteristics and growth rate between months, and one-way ANOVAs to test for differences among waterbody types. If differences were found from the ANOVA test, the Tukey HSD test was applied to identify which groups were different. These tests were performed using the *stats* package in R (CoreTeam, 2017). Four parameters (PP, NH_3_, N:P, and Chl-a) did not meet ANOVA criteria for equal variance, so the non-parametric Kruskal-Wallis and Kruskal-Wallis multiple comparison tests were applied to those parameters to test for differences among waterbody types.

To identify relationships between growth rates and site characteristics we applied a model selection approach using the Akaike Information Criterion (AIC). Simple linear regressions were used initially to determine which parameters should be included in the selection process, and multiple linear regressions were used for model selection. One outlier was removed from the blue-green growth rate data to evaluate regression models, and the mean growth rate from July and August experiments was used in the analyses. Non-normally distributed parameters were transformed when necessary. Unless otherwise noted, all analyses were conducted in R for Statistical Computing (V 3.5.0) using the *MASS* and *nlme* packages.

## 3. Results

### 3.1 Site characteristics

In 2017, water chemistry for composited samples was analyzed for each of the three zones (‘River’, ‘Lake’, and ‘Harbor’) (Table 3). The most notable difference in water chemistry among the zones was in NO_3_^−^ and Si, where the ‘River’ zone had much lower and higher concentrations than the ‘Lake’ and ‘Harbor’ zones, respectively. Also notable is the lower chl-a observed in the ‘River’ zone compared to the others. In 2018 water chemistry, temperature, and conductivity were measured at each site in July and August. We compared water quality data from each waterbody type (‘River’, ‘Lake/Pond’, and ‘Coastal’) using boxplots to illustrate differences among the three groups (Figure 2) and performed one-way ANOVAs with Tukey’s HSD test to determine differences in site characteristics and cyanobacterial growth rates among waterbody types (Table S1). Note that the ‘Coastal’ waterbody type refers to sites that are influenced by Lake Superior and inland rivers/streams. TP, TDP, and PP were similar among groups (Figure 2C, I, and F), as were particulate fractions of carbon and nitrogen (Figure 2D, E). TOC, DOC, TN, and TDN were lower in the ‘River’ than in the ‘Coastal’ and ‘Lake/Pond’ waterbody types (Figure 2A, G, B, and H), with statistical differences (p<0.05) between the ‘Lake/Pond’ and ‘River’ for TOC, DOC, and TDN. The N:P ratio was also lower in the ‘River’ group and had a much narrower range (Figure 2M), and NO_3_ ^−^ was highest in the ‘River’ group (Figure 2K).Chl-a was highest in the ‘Coastal’ waterbody type, following by ‘Lake/Pond’ and then ‘River’, with statistical differences between the ‘Coastal’ and ‘River’, and the ‘River’ and ‘Lake/Pond’ waterbody types. Temperature was lowest in the ‘River’ group and highest in the ‘Lake/Pond’ waterbodies (Figure 2O), with a statistical difference between the ‘Lake/Pond’ and ‘River’ waterbodies. EC_25_ was highest in the ‘Coastal’ waterbodies (Figure 2P), followed by the ‘River’ and then ‘Lake/Pond’. Paired two-sided t-tests were used to identify statistical differences in site characteristics and cyanobacterial growth rates between months. Significant (p<0.05) differences were observed for all parameters except for N:P, POC, PON, PP, chl-a, and growth rate (Table S2).

**Table 3.** Water chemistry means and standard deviations (analytical variation across a given sample) for composited water samples from each zone in 2017 experiments.

| Parameter | Lake |  | River |  | Harbor |  |
| --- | --- | --- | --- | --- | --- | --- |
|  | Mean | SD | Mean | SD | Mean | SD |
| Chl-a ($\mu\text{g/L}$) | 1.92 | 0.02 | 0.79 | 0.01 | 3.47 | 0.34 |
| POC ($\mu\text{M}$) | 26.49 | 1.37 | 43.30 | 1.19 | 41.11 | 3.17 |
| PON ($\mu\text{M}$) | 3.02 | 0.14 | 4.66 | 0.68 | 4.95 | 0.06 |
| NH<sub>3</sub> ($\mu\text{M}$) | 0.85 | 0.10 | 0.21 | 0.01 | 1.44 | 0.11 |
| NO<sub>3</sub><sup>-</sup> ($\mu\text{M}$) | 25.81 | - | 0.95 | - | 30.27 | - |
| Si ($\mu\text{M}$) | 25.42 | 0.04 | 121.45 | 0.71 | 50.53 | 0.02 |
| PP ($\mu\text{M}$) | 0.10 | 0.00 | 0.27 | 0.00 | 0.32 | 0.03 |
| TDP ($\mu\text{M}$) | 0.12 | - | 0.01 | - | 0.34 | - |
| SRP ($\mu\text{M}$) | 0.05 | - | 0.37 | - | 0.26 | - |
| TP ($\mu\text{M}$) | 0.16 | - | 0.52 | - | 0.76 | - |

**Figure 2.**
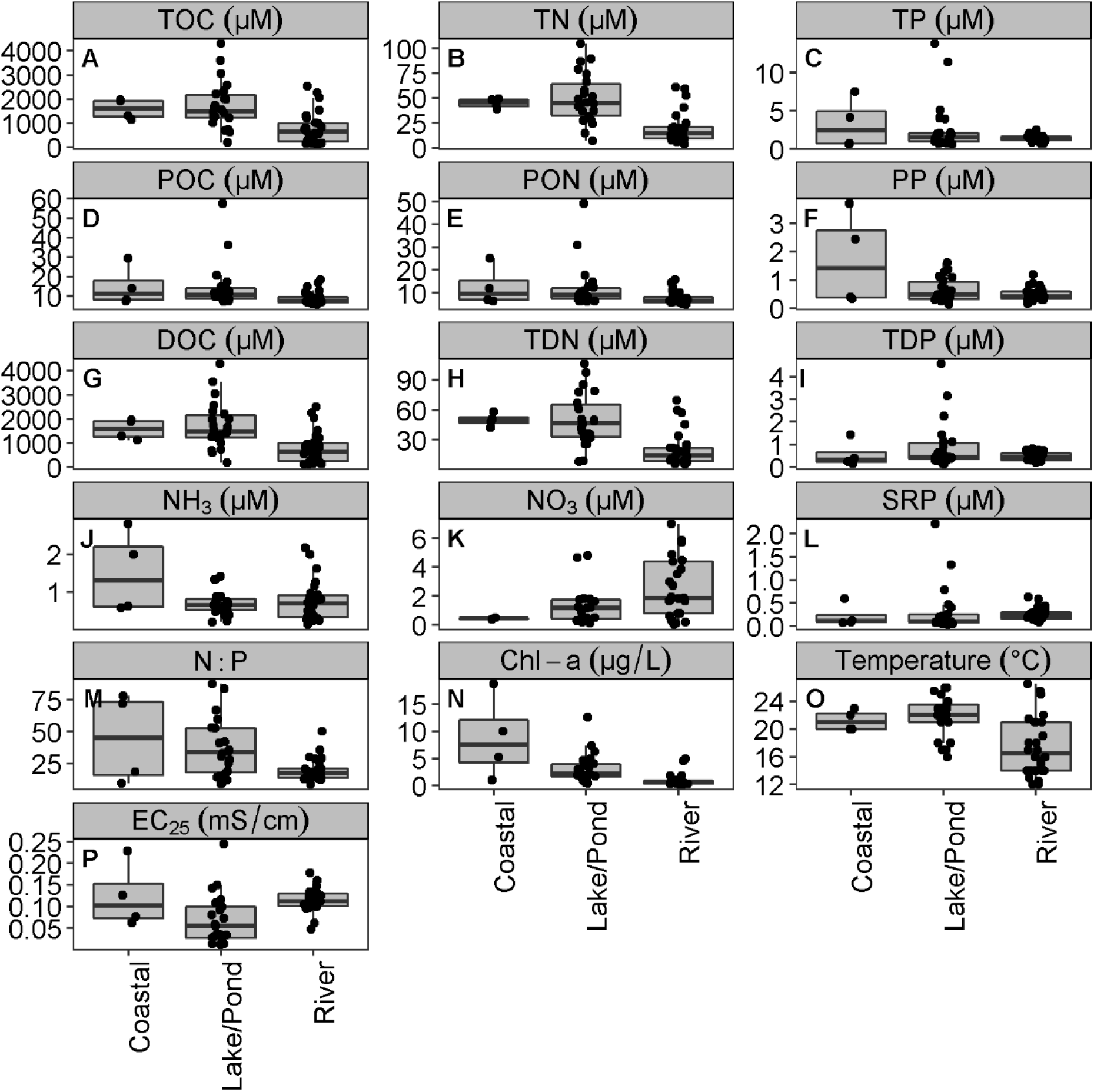
Boxplots of water quality parameters for each waterbody type in 2018 experiments. The central box line indicates the median, the upper and lower box hinges correspond to the 75% and 25% quartiles, respectively, and the whiskers correspond to 1.5 times the interquartile range. Black dots indicate sample points.

### 3.2 Experimental Results

The results of the 2017 experiments aimed at identifying zones that are sources of cyanobacteria showed that cyanobacteria grew fastest and achieved highest abundance in low N:P conditions at 20 and 25°C from the ‘River’ and ‘Harbor’ zones (Figure 3N, O, Q, and R, Table 4). Significant differences in main effects and all two-way interactions were observed for cyanobacterial growth (Table 4). Temperature alone had a significant effect on the “green” algal group, and all treatments and interactions had a significant effect on the “brown” algal group except for the interactions between N:P and temperature, and location and temperature.

**Table 4.** Results of the 3-way MANOVA test for each algal group using growth rate as the response variable. P-values marked with an asterisk (*) are significant at α=0.05).

| Treatment | Blue-green |  | Green |  | Brown |  |
| --- | --- | --- | --- | --- | --- | --- |
|  | F value | p-value | F value | p-value | F value | p-value |
| Temp | 6.09 | 0.0079<sup>*</sup> | 5.01 | 0.016<sup>*</sup> | 21.6 | 6.5 x 10<sup>-6*</sup> |
| Location | 7.71 | 0.0029<sup>*</sup> | 2.34 | 0.12 | 79.7 | 8.4 x 10<sup>-11*</sup> |
| N:P | 13.0 | 0.0015<sup>*</sup> | 0.940 | 0.34 | 66.2 | 4.5 x 10<sup>-8*</sup> |
| Location x Temp | 2.92 | 0.044<sup>*</sup> | 1.15 | 0.36 | 1.89 | 0.15 |
| N:P x Temp | 12.7 | 0.00022<sup>*</sup> | 0.380 | 0.69 | 2.38 | 0.12 |
| N:P x Location | 8.03 | 0.0024 | 0.290 | 0.75 | 18.1 | 2.2 x 10<sup>-5*</sup> |
| N:P x Location x Temp | 1.21 | 0.31 | 0.0260 | 0.97 | 6.10 | 0.0078<sup>*</sup> |

**Figure 3.**
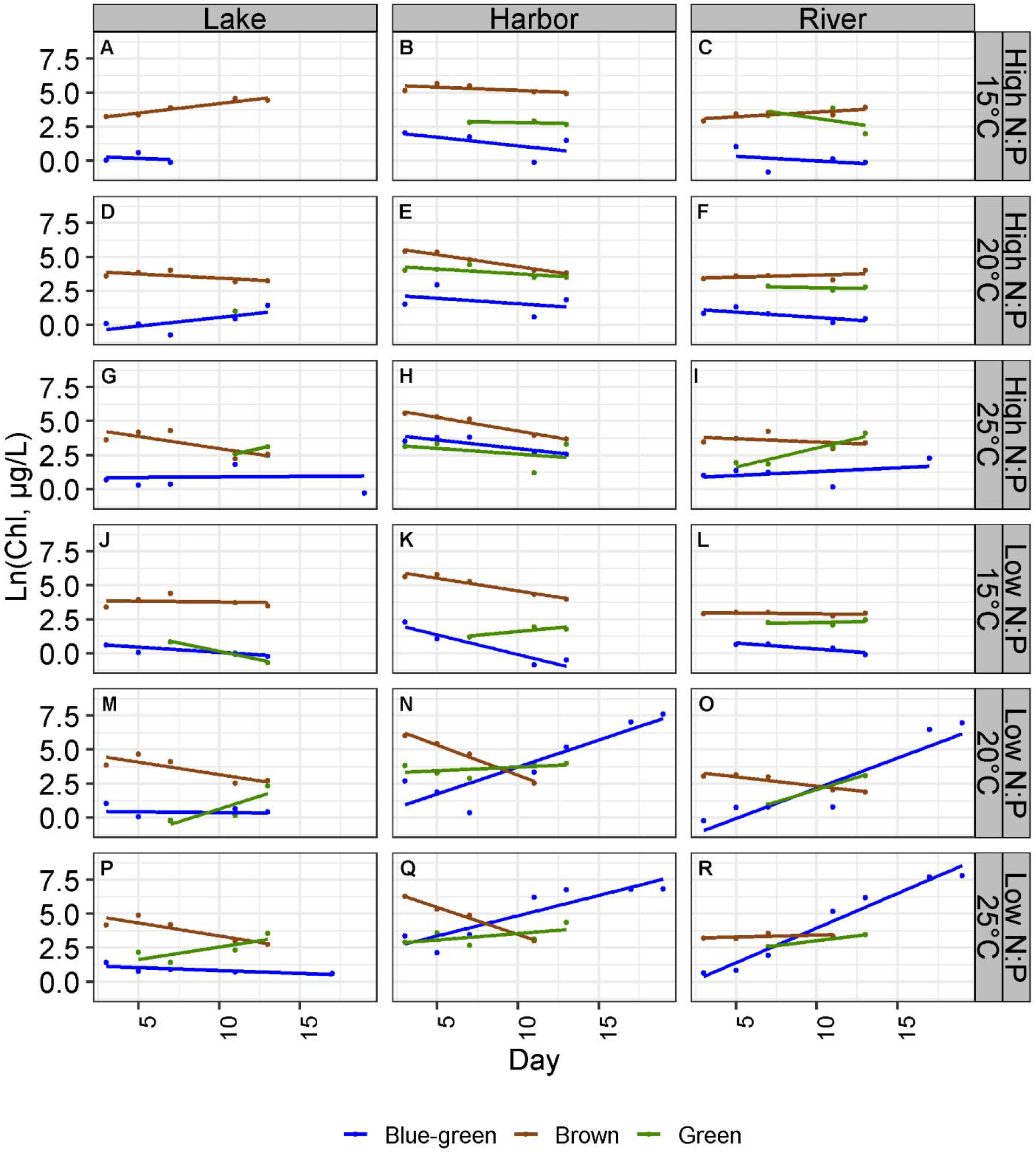
The natural log of chlorophyll concentration of the blue-green, green, and brown algal groups over the duration of the experiment for each treatment.

Taxonomic identification showed that cyanobacteria were present in cultures with high “blue-green” growth rates measured by the PHYTO-PAM, and that they were in highest abundance in the Low N:P and the 20°C and 25°C treatments (Figure 4). The results of the Kruskal Wallis tests (Table S3) agree with those found in the MANOVA, that there were significant differences in the relative abundance of *D. lemmermannii* between the high and low N:P treatments and the 20°C and 25°C temperature treatments; however, there were not always significant differences among locations for a given nutrient and temperature treatment combination. The relative abundance between the ‘River’ and ‘Harbor’ locations in the 20°C and low N:P treatment conditions were not different, and locations were not different in both temperature levels in the high N:P treatment with exception to 20°C-High-River and 25°C-High-Lake, where the abundance was negligible. The taxonomic results support those found using the PHYTO-PAM and also provide additional information: the species of cyanobacteria found in the ‘River’ and ‘Harbor’ locations is the same species observed in Lake Superior blooms, *D. lemmermannii,* and while cyanobacterial growth was high in both 20°C and 25°C at low N:P, *D. lemmermannii* specifically grew in the greatest relative abundance in 20°C and low N:P.

**Figure 4.**
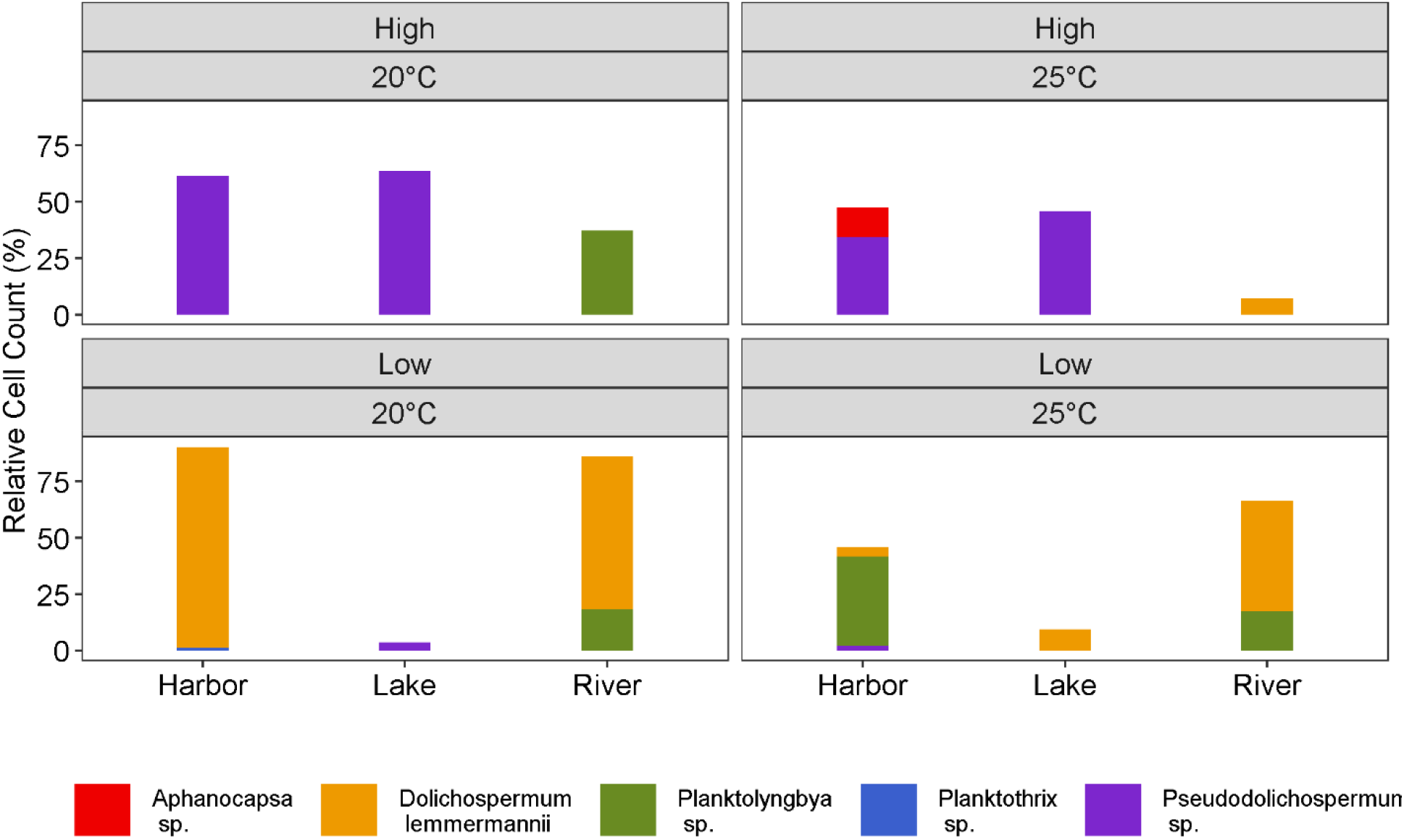
Relative abundance of cyanobacterial species by treatment combinations. The 15°C treatment was not analyzed due to mechanical failure of the incubator.

The results from the 2017 experiments indicated that upland waters contribute to cyanobacterial blooms in the lake, so in 2018 we evaluated 26 specific locations (rivers, lakes/ponds, and coastal waterbodies) as potential sources of cyanobacterial propagules and looked for characteristics shared among sites that could be potential sources. The results from 2018 showed that sites with the highest cyanobacterial growth rates were widely scattered across the study region (Figure 5). There were no spatial autocorrelations in the growth data as represented in Moran’s I. We then looked for similarities in chemical, biological, and physical characteristics that could identify sites with high potential using single and multiple linear regressions.

**Figure 5.**
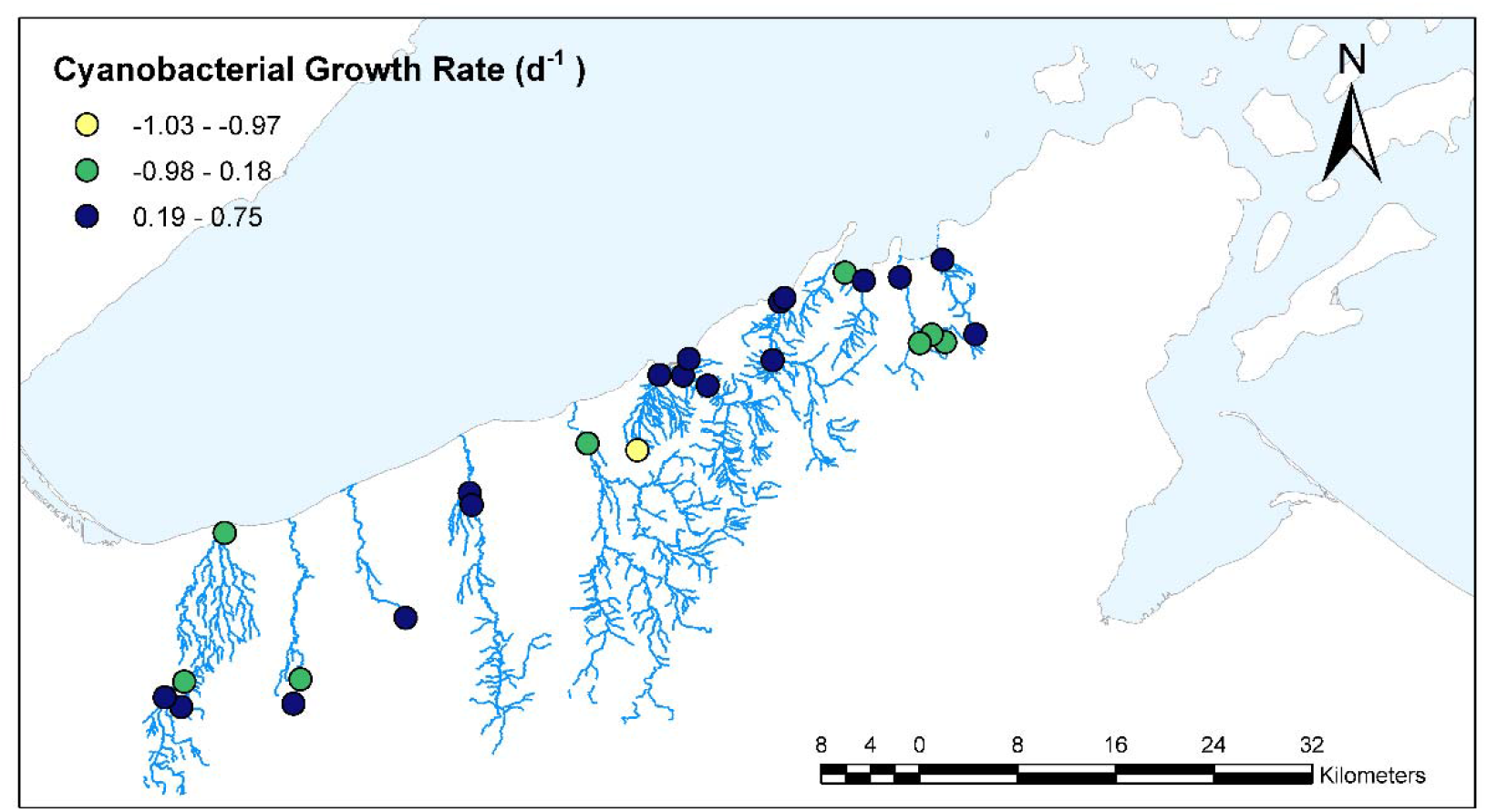
Sampling locations of experimental cyanobacterial growth rate for the months of July and August. Growth rates are divided by natural breaks.

Simple linear regressions were used to compare cyanobacterial growth rates to site characteristics. Out of 17 water quality parameters tested, four were significant predictors for cyanobacterial growth: temperature (p = 0.006, R^2^ = 0.26, slope = −0.026), EC_25_ (p = 0.0004, R^2^ = 0.41, slope = 2.70), ln(SRP) (p = 0.002, R^2^ = 0.34, slope = 0.17), and ln(N:P) (p = 0.02, R^2^ = 0.20, slope = −0.17). These results mean that cyanobacteria grew fastest under laboratory conditions from sites that had higher conductivity and SRP and lower temperature and N:P. Those parameters were then used with AIC and backward model selection to identify the best model for cyanobacterial growth rate. The optimized model included only *in situ* EC_25_ and temperature (Table 5). Model selection could only be performed using additive models (i.e. without interactions), as the number of comparisons with multiplicative models would be too l computationally expensive, therefore we tested the best additive model for interaction terms and found significant interaction between temperature and EC_25_, which resulted in a final model with R^2^ = 0.57 (Table 6). We also tested for the impact of waterbody type (Coastal, River, Lake/Pond) in the model and found that there was no significant effect.

**Table 5.** Results of model selection using AIC.

| Model (N=24) | AIC |
| --- | --- |
| Ln(SRP) + Ln(NP) + Temp + EC<sub>25</sub> | -85.45 |
| Ln(SRP) + Temp + EC<sub>25</sub> | -87.41 |
| Temp + EC<sub>25</sub> | -88.58 |
| EC<sub>25</sub> | -88.84 |

**Table 6.** Summary of interaction model using temperature and EC_25_.

| Parameter | Slope Estimate | p-value | Overall<br>R<sup>2</sup> | Overall<br>p-value |
| --- | --- | --- | --- | --- |
| EC<sub>25</sub> | 15.7 | 0.00384 | 0.57 | 0.00015 |
| Temp | 0.0542 | 0.0384 |  |  |
| Temp x EC<sub>25</sub> | 0.227 | 0.00990 |  |  |

To validate measurements using the PHYTO-PAM, we also examined preserved samples for identification. Preserved samples from 14 sites with the highest final “blue-green” biomass (measured using the PHTYO-PAM) were surveyed. This group included one ‘Coastal’ waterbody, four ‘Lake/Pond’, and nine ‘River’ sites. Seven of these samples were found to be comprised of over 75% *D. lemmermannii* and three sites had greater than 50% (Figure 6). Ten out of the total 14 sites were also among sites with the highest experimental growth rates for “blue-green” algae (calculated using PHYTO-PAM data). Three out of the four sites that were not among the highest observed growth rates were ‘Lake/Pond’ waterbody types, and two of those four sites were comprised of less than 25% *D. lemmermannii* (1 ‘River’ and 1 ‘Lake/Pond’).

**Figure 6.**
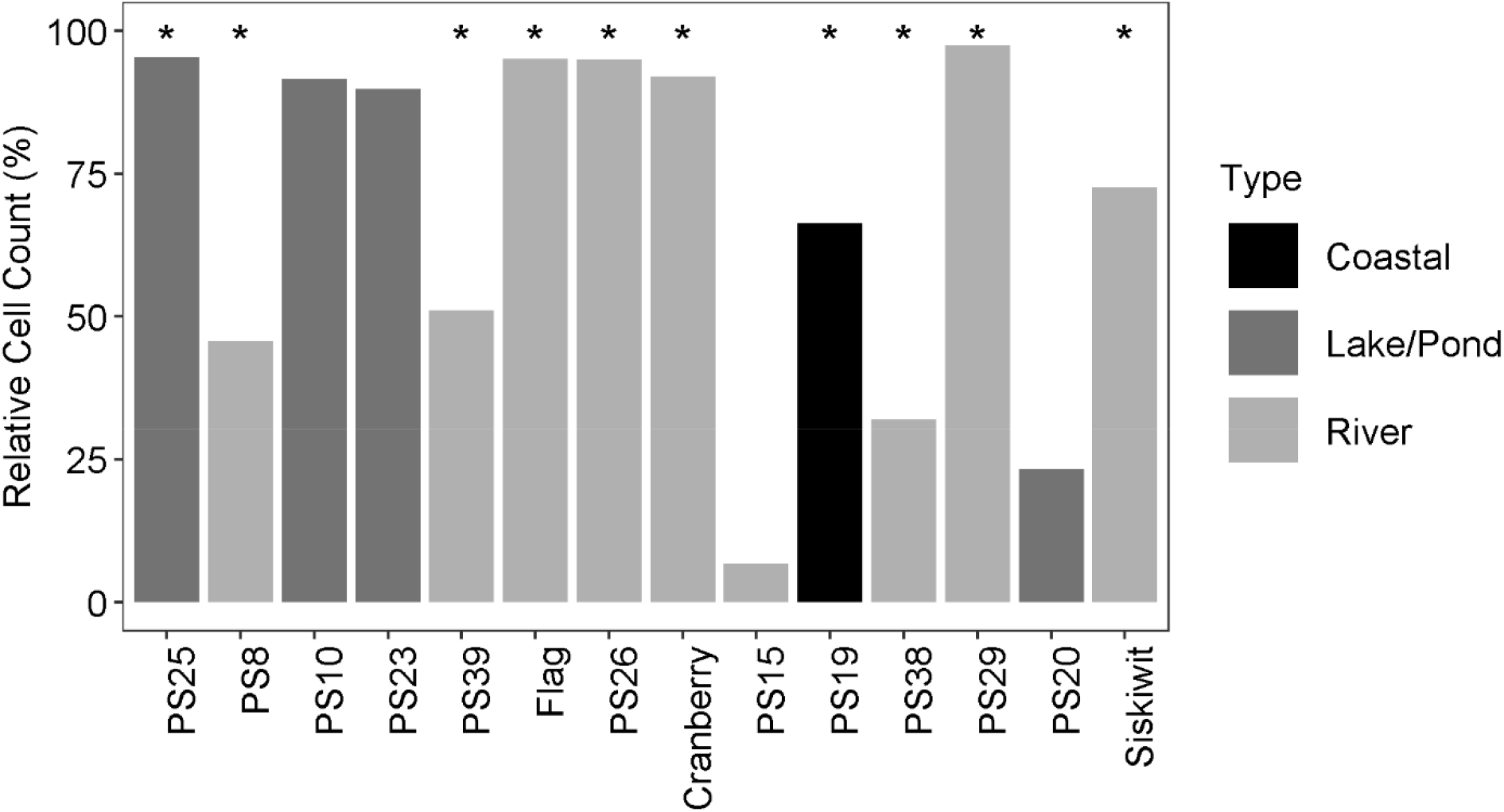
Relative abundance of *D. lemmermannii* in 14 sites (ordered from west to east) with greatest cyanobacterial growth in culture. Colors indicate waterbody type and asterisks (*) denote the 10 sites that also had the highest cyanobacterial growth rates.

## 4 Discussion

The appearance of cyanobacterial blooms in Lake Superior is surprising, given that it is a cold, oligotrophic system with much less anthropogenic nutrient inputs relative to other lakes where blooms are often observed. The current “chemostat paradigm” for cyanobacterial blooms in lakes is that the role of rivers is to supply nutrients to algal populations in the receiving lakes (Bridgeman et al., 2012; Carpenter et al., 1998; Conroy, 2007), but this study points to other potentially important roles for inflowing waters. This work is a critical step toward understanding why we have begun observing cyanobacterial blooms in Lake Superior and it indicates a likely important role of upland waterbodies, particularly rivers, in seeding blooms. There is a wealth of evidence in marine and freshwater systems that bloom-forming cyanobacteria can persist in rivers and estuaries (Conroy et al., 2014; Paerl and Otten, 2013; Reif, 1939; Schwartz, 2007; Steinberg and Hartman, 1988). What is not as well understood is how the presence of cyanobacteria in rivers may affect downstream lakes or ponds. The lack of research on this topic may be due to the greater likelihood of observing cyanobacterial blooms in lentic environments than in their lotic counterparts or because riverine conditions (including high turbulence, low light, short residence times, and cooler temperatures) are not typically associated with high algal growth rates (Paerl and Otten, 2013). Nevertheless, a recent study of 11 major rivers in the US showed that cyanobacteria was present in all rivers and comprised up to 52% of the community composition (Graham et al., 2020). The present study shows that rivers may play a critical role in the promotion of newly occurring cyanobacterial blooms in Lake Superior.

Our work is based on the logic that if an environmental sample can generate cyanobacterial growth under reasonable time scales and favorable conditions in a laboratory setting, it has elevated potential to be a source of blooms compared to samples that do not produce high growth in the lab. This approach has the advantage of not relying on quantifying potentially rare populations that initiate blooms, and it avoids the potential that a given algal taxon may be present but not be viable, issues that could arise under a simple sampling regime no matter how sensitive it may be. The experiments we conducted in 2017 and 2018 provided insight into the origins and the conditions that promote cyanobacterial blooms along south shore of Lake Superior. The conclusions we draw from the 2017 experiment are that bloom-forming cyanobacteria grew abundantly in low N:P and at moderate to high temperature, and more importantly, only from ‘River’ and ‘Harbor’ locations and not from the ‘Lake’ samples. This finding provides evidence that blooms are unlikely to have been sourced just from the lake, and that upstream sources are likely important contributors of living cells to generate blooms. The subsequent 2018 work then began to narrow down characteristics of upstream environments that are most likely to be involved in the generation and delivery of algal seed populations for Lake Superior blooms.

Cyanobacteria in the Nostocales order are diazotrophic; thus, it is not surprising that *D. lemmermannii* grew better than the other algal groups in these nitrogen-limiting conditions during laboratory experiments, but growth under lab conditions does not necessarily indicate the conditions most responsible for high growth in a natural setting. PHYTO-PAM measurements in both 2017 and 2018 did not show a dominance of cyanobacteria in river samples at the beginning of the experiments. In fact, the initial total biomass of samples was low and typically dominated by the “brown” algal group while the “blue-green” only comprised approximately 1-5% of the total biomass (Figure 3). This was also true for ‘Lake’ and ‘Harbor’ samples from 2017, which may mean that while similar concentrations of cyanobacteria were initially present from all zones, cells from ‘River’ and ‘Harbor’ samples were more likely *D. lemmermannii* and samples from the ‘Lake’ sites were not, given that this species did not grow in high abundance in any treatment combination from this location. In an analysis of Lake Superior nearshore and offshore phytoplankton survey data from 2011 and 2016, (Kovalenko et al., 2019) found that diatoms accounted for over 75% of the total biovolume for all samples areas, with some codominance in 2016. Codominance of *D. lemmermannii* (by biovolume) was observed for part of 2016 at one site in central Lake Superior offshore (depth > 200m), indicating that *D. lemmermannii* is not prominent in the Lake Superior phytoplankton community. ‘Harbor’ samples did grow cyanobacteria in abundance, but a closer look at the taxonomic data showed that the species present from ‘Harbor’ samples was not predominately *D. lemmermannii* with exception to the low N:P and 20 °C treatment. Analysis of 2018 data revealed that ‘River’ sites had the lowest total N:P ratio of the three waterbody types (mean of approximately19:1); therefore, from a N:P standpoint, rivers feeding into nearshore Lake Superior where blooms have been observed may provide conditions to promote diazotrophic cyanobacteria like *D. lemmermannii*. Consequently, even though the relative abundance of cyanobacteria is not greatest in the overall community composition, the conditions in rivers support a community of cyanobacteria that is capable of proliferating under laboratory conditions.

In addition to a clear N:P preference, higher water temperatures stimulated growth during the 2017 experiments, and more specifically, taxonomic data showed that the 20°C treatment resulted in higher *D. lemmermannii* abundances than 25°C. We also observed high cyanobacterial growth rates in the 2018 experiments under laboratory conditions (25°C and low N:P), but the sites that produced those high growth rates under laboratory conditions, namely rivers, were characterized by low water temperatures *in situ* (12-26°C, mean of approx. 17 °C). These findings indicate that fast-growing cyanobacteria originated in colder waters, but our 2017 experiments showed that 15°C was too cold for extensive growth of *D. lemmermannii*, even at optimal nutrient conditions. Evidence from subalpine lakes has shown the ability of *D. lemmermannii* to persist in cold water temperature systems and then grow extensively as temperatures warm to greater than 15°C (Salmaso et al., 2015), which is consistent with the temperature dependence we observed in our study.

Routine water quality monitoring during the 2018 field season (May-October) showed that the lake is typically warmer than inflowing rivers by several degrees (Figure 7). Cooler river temperatures can be attributed to several factors including size, flowrate, canopy cover, hydrologic inputs, and drainage pattern, whereas water in shallow nearshore embayments is exposed to direct light and the surface layer warms throughout the summer, resulting in higher surface water temperatures in the nearshore than in rivers. Temperature has been identified as a primary control on *D. lemmermannii* growth in other cold, oligotrophic lakes (Callieri et al., 2014; Capelli et al., 2017; Salmaso et al., 2015, 2012), and has also been identified as an important driver in Lake Superior (Sterner et al., in review). So, while rivers are more likely to yield viable *D. lemmermannii* cells that could initiate growth in the lake, the warmer temperature conditions in Lake Superior’s nearshore are generally more favorable for growing high concentrations of biomass. Light conditions in the nearshore and rivers was not measured during routine monitoring; however, river plumes in Lake Superior have been studied and it has been shown that light is lower in Lake Superior waters impacted by river plumes (Cooney et al., 2018; Minor et al., 2014), indicating that rivers have higher turbidity and thus, lower light compared to the nearshore. Detenbeck et al. (2004) also document high turbidity in south shore streams due to forest type and erosion of fine sediment with mean field turbidity measurements near approximately 100 NTU, which corresponds to an extinction coefficient of 5 m^−1^ (Brown, 1984), while typical Lake Superior values are less than approximately 0.4 m^−1^ (Cooney et al., 2018; Sterner, 2010). We hypothesize that when *D. lemmermannii* propagules from rivers with low temperature and light conditions reach the nearshore of Lake Superior, where temperature is seasonally warmer and there is greater light exposure, they can grow in greater abundance and ultimately produce blooms. Further work is needed, however, to understand the relative contributions of temperature, light, and nutrients, as well as hydrodynamics, in promoting cyanobacterial blooms once the cells have entered the lake. Additional work is also needed to determine whether rivers are supporting a community of *D. lemmermannii* or acting as a transport mechanism from other upstream areas.

**Figure 7.**
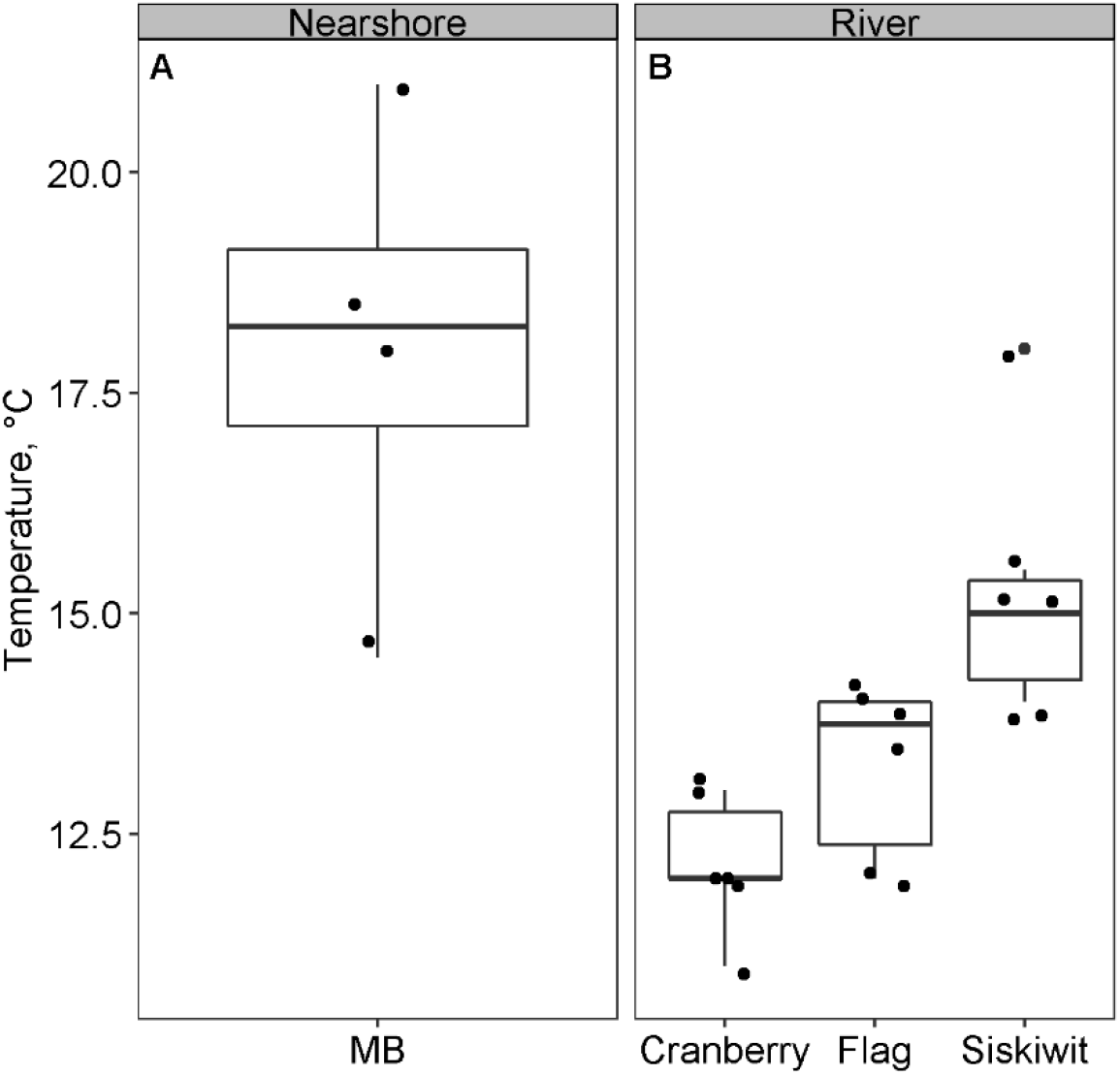
Boxplots of July and August surface water temperature for nearshore Lake Superior (MB) and a subset of rivers included in 2018 experiments. The central box line indicates the median, the upper and lower box hinges correspond to the 75% and 25% quartiles, respectively, and the whiskers correspond to 1.5 times the interquartile range. Black dots indicate sample points.

The other predictor that helped explain which sites are likely sources of cyanobacterial propagules to Lake Superior, in addition to colder water temperatures, was higher conductivity. Conductivity is related to salinity and pH, both of which can affect cyanobacteria growth. There is some evidence that salinity can inhibit growth (Apte et al., 1987; Rai, 1990), but the concentrations used in these studies (55 mM – 500 mM NaCl) typically far exceed those we observed in our study sites (approximately 2.2 mM NaCl, assuming that all of the conductivity can be attributed to salinity). Similarly, pH conditions in five south shore Lake Superior streams were circumneutral during multiple sampling events in 2016, indicating that pH is not likely a factor influencing algal growth (J. Delvaux, unpublished data). Hence, it is unlikely that the levels of conductivity we observed are high enough to have an inhibitory effect on growth, and the more likely explanation is that conductivity is related to some other unmeasured parameter that is related to sites with high potential. Conductivity may be related to the geological substrates in the region. The predominate soil types in the study area are Alfisols and Spodosols, commonly referred to as ‘red clay’ due to their rich iron content, are highly susceptible to erosion and the main contributor of dissolved ions to Lake Superior (Cooney et al., 2018; Stortz et al., 1976). We considered the high iron content of the soil as a possible linkage between high potential sites and increased conductivity, but the iron in these soils is present as iron oxides and are highly insoluble in water, and so is not likely stimulating algal growth (Tonello et al., 2019).

To summarize, evidence from this study shows that rivers play a critical role in initiating cyanobacterial bloom on the south shore of Lake Superior, with some additional evidence that shifting growth conditions between rivers and lakes are also necessary for bloom formation. Our data showed that *D. lemmermannii* grew under laboratory conditions from water samples that were collected from inland locations, particularly rivers, and grew best in 20°C and low N:P conditions. Somewhat unexpectedly, sites that had the greatest cyanobacterial growth under optimal laboratory conditions were characterized by relatively low temperatures and high conductivity. We ascertain that rivers produce cells of *D. lemmermannii* capable of initiating extensive growth, but cold water temperatures and low light conditions inhibit overall growth in these habitats. When these cells enter the Lake Superior nearshore, where temperatures are seasonally warmer and light levels are higher, conditions are more favorable for growth.

Based on this evidence, we argue that rivers provide a consistent source of viable propagules of *D. lemmermannii* to Lake Superior, and that the appearance of blooms in the lake is dependent on in-lake conditions. Lake Superior has already begun experiencing significant warming (Austin and Colman, 2007) and extreme precipitation events (Cooney et al., 2018), and these trends are expected to continue. If water temperatures and nutrient inputs continue to increase, along with consistent seeding from rivers, cyanobacterial blooms may become a persistent feature of Lake Superior’s nearshore.

## Author Contribution Statement

All authors have read and approved the final submission. KLR developed experimental design, carried out laboratory and field research, conducted data analysis, and prepared this manuscript. RWS advised on experimental design and data analysis, and provided significant comments on the manuscript. BLM advised on experimental design, assisted with data collection, and commented on data analyses and manuscript. SLB assisted with field and laboratory research, sample analysis, and commented on the manuscript.

## Acknowledgments

We would like to acknowledge and thank Nicole Farley and Madison Perry for their assistance in the lab and in the field. Their tireless efforts and positive attitudes made this project possible and enjoyable.

## References

Anderson, D.M., 1989. Toxic Algal Blooms and Red tides: a Global Perspective. Biol. Environ. Sci. Toxicol. https://doi.org/10.1029/95rg00440

Apte, S.K., Reddy, B.R., Thomas, J., 1987. Relationship between Sodium Influx and Salt Tolerance of Nitrogen-Fixing Cyanobacteria. Appl. Environ. Microbiol. 53, 1934–9.

Atkins, P., de Paula, J., 2006. Physical Chemistry, 8th ed. W. H. Freeman and Company, New York.

Austin, J.A., Colman, S.M., 2007. Lake Superior summer water temperatures are increasing more rapidly than regional temperatures: A positive ice-albedo feedback. Geophys. Res. Lett. 34, L06604. https://doi.org/10.1029/2006GL029021

Baer, K.N., Goulden, C.E., 1998. Evaluation of a high-hardness COMBO medium and frozen algae for Daphnia magna. Ecotoxicol. Environ. Saf. 39, 201–206. https://doi.org/10.1006/eesa.1997.1627

Bridgeman, T.B., Chaffin, J.D., Kane, D.D., Conroy, J.D., Panek, S.E., Armenio, P.M., 2012. From River to Lake: Phosphorus partitioning and algal community compositional changes in Western Lake Erie. J. Great Lakes Res. 38, 90–97. https://doi.org/10.1016/j.jglr.2011.09.010

Brown, R., 1984. Relationships between suspended solids, turbidity, light attenuation, and algal productivity. Lake Reserv. Manag. 1, 198–205. https://doi.org/10.1080/07438148409354510

Callieri, C., Bertoni, R., Contesini, M., Bertoni, F., 2014. Lake Level Fluctuations Boost Toxic Cyanobacterial ‘’Oligotrophic Blooms. PLoS One 9, 109526. https://doi.org/10.1371/journal.pone.0109526

Capelli, C., Ballot, A., Cerasino, L., Papini, A., Salmaso, N., 2017. Biogeography of bloom-forming microcystin producing and non-toxigenic populations of Dolichospermum lemmermannii (Cyanobacteria). Harmful Algae 67, 1–12. https://doi.org/10.1016/j.hal.2017.05.004

Carpenter, S.R., Caraco, N.F., Correll, D.L., Howarth, R.W., Sharpley, A.N., Smith, V.H., 1998. Nonpoint pollution of surface waters with phosphorus and nitrogen. Ecol. Appl. 8, 559–568. https://doi.org/10.1890/1051-0761(1998)008[0559:NPOSWW]2.0.CO;2

Chaffin, J.J.D., Sigler, V., Bridgeman, T.B.T., Justin D. Chaffin1, V.S., Thomas B. Bridgeman, 2014. Connecting the blooms: Tracking and establishing the origin of the record-breaking Lake Erie Microcystis bloom of 2011 using DGGE. Aquat. Microb. Ecol. 73, 29–39. https://doi.org/10.3354/ame01708

Chapra, S.C., Dove, A., Warren, G.J., 2012. Long-term trends of Great Lakes major ion chemistry. J. Great Lakes Res. 38, 550–560. https://doi.org/10.1016/j.jglr.2012.06.010

Cloern, J.E., 1996. Phytoplankton bloom dynamics in coastal ecosystems: A review with some general lessons from sustained investigation of San Francisco Bay, California. Rev. Geophys. https://doi.org/10.1029/96RG00986

Cloern, J.E., Jassby, A.D., 2010. Patterns and Scales of Phytoplankton Variability in Estuarine–Coastal Ecosystems. Estuaries and Coasts 33, 230–241. https://doi.org/10.1007/s12237-009-9195-3

Cloern, J.E., Jassby, A.D., 2008. Complex seasonal patterns of primary producers at the land-sea interface. Ecol. Lett. 11, 1294–1303. https://doi.org/10.1111/j.1461-0248.2008.01244.x

Conroy, J.D., 2007. Testing the algal loading hypothesis: the importance of Sandusky River phytoplankton inputs to offshore Lake Erie processes. Diss. Abstr. Int.

Conroy, J.D., Culver, D.A., Heath, R.T., 2008. Gloom and blooms: simulating phytoplankton growth moving out of a tributary into a large lake. SIL Proceedings, 1922-2010 30, 615–618. https://doi.org/10.1080/03680770.2008.11902201

Conroy, J.D., Kane, D.D., Briland, R.D., Culver, D.A., 2014. Systemic, early-season Microcystis blooms in western Lake Erie and two of its major agricultural tributaries (Maumee and Sandusky rivers). J. Great Lakes Res. 40, 518–523. https://doi.org/10.1016/j.jglr.2014.04.015

Cooney, E.M., McKinney, P., Sterner, R., Small, G.E., Minor, E.C., 2018. Tale of Two Storms: Impact of Extreme Rain Events on the Biogeochemistry of Lake Superior. J. Geophys. Res. Biogeosciences 123, 1719–1731. https://doi.org/10.1029/2017JG004216

CoreTeam, R., 2017. R: A Language and Environment for Statistical Computing.

Davis, T.W., Watson, S.B., Rozmarynowycz, M.J., Ciborowski, J.J.H., Mckay, R.M., Bullerjahn, G.S., 2014. Phylogenies of microcystin-Producing cyanobacteria in the lower laurentian great lakes suggest extensive genetic connectivity. PLoS One 9, e106093. https://doi.org/10.1371/journal.pone.0106093

Detenbeck, N.E., Elonen, C.M., Taylor, D.L., Anderson, L.E., Jicha, T.M., Batterman, S.L., 2004. Region, landscape, and scale effects on Lake Superior tributary water quality. J. Am. Water Resour. Assoc. 40, 705–720. https://doi.org/10.1111/j.1752-1688.2004.tb04454.x

Graham, J.L., Dubrovsky, N.M., Foster, G.M., King, L.R., Loftin, K.A., Rosen, B.H., Stelzer, E.A., 2020. Cyanotoxin occurrence in large rivers of the United States. Inl. Waters 1–9. https://doi.org/10.1080/20442041.2019.1700749

Guillard, R.R.L., 1975. Culture of Phytoplankton for Feeding Marine Invertebrates, in: Culture of Marine Invertebrate Animals. Springer US, Boston, MA, pp. 29–60. https://doi.org/10.1007/978-1-4615-8714-9_3

Holmes, R.M., Aminot, A., Kérouel, R., Hooker, B.A., Peterson, B.J., 1999. A simple and precise method for measuring ammonium in marine and freshwater ecosystems. Can. J. Fish. Aquat. Sci. 56, 1801–1808. https://doi.org/10.1139/f99-128

Howell, E.T., Chomicki, K.M., Kaltenecker, G., 2012. Tributary discharge, lake circulation and lake biology as drivers of water quality in the Canadian Nearshore of Lake Ontario. J. Great Lakes Res. 38, 47–61. https://doi.org/10.1016/j.jglr.2012.03.008

Huisman, J., Codd, G.A., Paerl, H.W., Ibelings, B.W., Verspagen, J.M.H., Visser, P.M., 2018. Cyanobacterial blooms. Nat. Rev. Microbiol. https://doi.org/10.1038/s41579-018-0040-1

Kilham, S.S., Kreeger, D.A., Lynn, S.G., Goulden, C.E., Herrera, L., 1998. COMBO: A defined freshwater culture medium for algae and zooplankton. Hydrobiologia 377, 147–159. https://doi.org/10.1023/A:1003231628456

Kling, G.W., Kipphut, G.W., Miller, M.M., O’Brien, W.J., 2000. Integration of lakes and streams in a landscape perspective: The importance of material processing on spatial patterns and temporal coherence. Freshw. Biol. 43, 477–497. https://doi.org/10.1046/j.1365-2427.2000.00515.x

Konopka, A., Brock, T.D., 1978. Effect of Temperature on Blue-Green Algae (Cyanobacteria) in Lake Mendota. Appl. Environ. Microbiol. 36, 572–576.

Kosten, S., Huszar, V.L.M., Bécares, E., Costa, L.S., van Donk, E., Hansson, L.A., Jeppesen, E., Kruk, C., Lacerot, G., Mazzeo, N., De Meester, L., Moss, B., Lürling, M., Nõges, T., Romo, S., Scheffer, M., 2012. Warmer climates boost cyanobacterial dominance in shallow lakes. Glob. Chang. Biol. 18, 118–126. https://doi.org/10.1111/j.1365-2486.2011.02488.x

Kovalenko, K.E., Reavie, E.D., Bramburger, A.J., Cotter, A., Sierszen, M.E., 2019. Nearshore-offshore trends in Lake Superior phytoplankton. J. Great Lakes Res. 45, 1197–1204. https://doi.org/10.1016/j.jglr.2019.09.016

Kratz, T.K., Webster, K.E., Bowser, C.J., Magnuson, J.J., Benson, B.J., 1997. The influence of landscape position on lakes in northern Wisconsin. Freshw. Biol. https://doi.org/10.1046/j.1365-2427.1997.00149.x

Kutovaya, O.A., McKay, R.M.L., Beall, B.F.N., Wilhelm, S.W., Kane, D.D., Chaffin, J.D., Bridgeman, T.B., Bullerjahn, G.S., 2012. Evidence against fluvial seeding of recurrent toxic blooms of Microcystis spp. in Lake Erie’s western basin. Harmful Algae 15, 71–77. https://doi.org/10.1016/j.hal.2011.11.007

Loftin, K.A., Clark, J.M., Journey, C.A., Kolpin, D.W., Van Metre, P.C., Carlisle, D., Bradley, P.M., 2016. Spatial and temporal variation in microcystin occurrence in wadeable streams in the southeastern United States. Environ. Toxicol. Chem. 35, 2281–2287. https://doi.org/10.1002/etc.3391

Lürling, M., Mello, M.M., van Oosterhout, F., Domis, L. de S., Marinho, M.M., 2018. Response of natural cyanobacteria and algae assemblages to a nutrient pulse and elevated temperature. Front. Microbiol. 9, 1851. https://doi.org/10.3389/fmicb.2018.01851

Minor, E.C., Forsman, B., Guildford, S.J., 2014. The effect of a flood pulse on the water column of western Lake Superior, USA. J. Great Lakes Res. 40, 455–462. https://doi.org/10.1016/j.jglr.2014.03.015

Moran, P.A.P., 1950. Notes on Continuous Stochastic Phenomena. Biometrika 37, 17. https://doi.org/10.2307/2332142

Murphy, J., Riley, J.P., 1962. A modified single solution method for the determination of phosphate in natural waters. Anal. Chim. Acta 27, 31–36. https://doi.org/10.1016/S0003-2670(00)88444-5

Paerl, H.W., Huisman, J., 2009. Climate change: A catalyst for global expansion of harmful cyanobacterial blooms. Environ. Microbiol. Rep. 1, 27–37. https://doi.org/10.1111/j.1758-2229.2008.00004.x

Paerl, H.W., Huisman, J., 2008. Climate: Blooms like it hot. Science (80-.). https://doi.org/10.1126/science.1155398

Paerl, H.W., Otten, T.G., 2013. Harmful Cyanobacterial Blooms: Causes, Consequences, and Controls. Microb. Ecol. 65, 995–1010. https://doi.org/10.1007/s00248-012-0159-y

Rai, A.K., 1990. Biochemical characteristics of photosynthetic response to various external salanities in halotolerant and fresh water cyanobacteria. FEMS Microbiol. Lett. 69, 177–180. https://doi.org/10.1111/j.1574-6968.1990.tb04196.x

Reif, C.B., 1939. The Effect of Stream Conditions on Lake Plankton. Trans. Am. Microsc. Soc. 58, 398. https://doi.org/10.2307/3222782

Roache, A., Cousins, R.M., Muszynski, M.R., 2020. Evaluation of the 2018 “Father’s Day Flood” Using Technology-Based Tools, in: Geotechnical Special Publication. American Society of Civil Engineers (ASCE), pp. 788–798. https://doi.org/10.1061/9780784482797.077

Robarts, R.D., Zohary, T., 1987. Temperature effects on photosynthetic capacity, respiration, and growth rates of bloomLforming cyanobacteria. New Zeal. J. Mar. Freshw. Res. 21, 391–399. https://doi.org/10.1080/00288330.1987.9516235

Salmaso, N., Buzzi, F., Garibaldi, L., Morabito, G., Simona, M., Letizia, @bullet, @bullet, G., Morabito, G., Simona, M., 2012. Effects of nutrient availability and temperature on phytoplankton development: a case study from large lakes south of the Alps. Aquat. Sci. 74, 555–570. https://doi.org/10.1007/s00027-012-0248-5

Salmaso, N., Capelli, C., Shams, S., Cerasino, L., 2015. Expansion of bloom-forming Dolichospermum lemmermannii (Nostocales, Cyanobacteria) to the deep lakes south of the Alps: Colonization patterns, driving forces and implications for water use. Harmful Algae 50, 76–87. https://doi.org/10.1016/J.HAL.2015.09.008

Schindler, D.W., 1975. Whole-lake eutrophication experiments with phosphorus, nitrogen and carbon. SIL Proceedings, 1922-2010 19, 3221–3231. https://doi.org/10.1080/03680770.1974.11896436

Schwartz, W., 2007. G. Evelyn Hutchinson, A Treatise on Limnology, Vol. II. Introduction to Lake Biology and the Limnoplankton. IX und 1115 S., 253 Abb., 53 Tab., 1 Taf. New York, London, Sydney 1967: John Wiley and Sons s 310.-. Z. Allg. Mikrobiol. 8, 477–477. https://doi.org/10.1002/jobm.19680080520

Siegel, S., Castellan, N.J., 1988. Nonparametric Statistics for the Behavioral Sciences, 2nd ed, The American Catholic Sociological Review. McGraw-Hill, New York. https://doi.org/10.2307/3708383

Steinberg, C.E.W., Hartman, H.M., 1988. Planktonic bloomLforming Cyanobacteria and the eutrophication of lakes and rivers. Freshw. Biol. 20, 279–287. https://doi.org/10.1111/j.1365-2427.1988.tb00452.x

Sterner, R.W., 2010. In situ-measured primary production in Lake Superior. J. Great Lakes Res. 36, 139–149. https://doi.org/10.1016/J.JGLR.2009.12.007

Stortz, K., Clapper, R., Sydor, M., 1976. Turbidity Sources in Lake Superior. J. Great Lakes Res. 2, 393–401. https://doi.org/10.1016/S0380-1330(76)72302-5

Sukenik, A., Quesada, A., Salmaso, N., 2015. Global expansion of toxic and non-toxic cyanobacteria: effect on ecosystem functioning. Biodivers. Conserv. 24, 889–908. https://doi.org/10.1007/s10531-015-0905-9

Taranu, Z.E., Gregory-Eaves, I., Leavitt, P.R., Bunting, L., Buchaca, T., Catalan, J., Domaizon, I., Guilizzoni, P., Lami, A., Mcgowan, S., Moorhouse, H., Morabito, G., Pick, F.R., Stevenson, M.A., Thompson, P.L., Vinebrooke, R.D., 2015. Acceleration of cyanobacterial dominance in north temperate-subarctic lakes during the Anthropocene. Ecol. Lett. 18, 375–384. https://doi.org/10.1111/ele.12420

Taylor, B.W., Keep, C.F., Hall, R.O., Koch, B.J., Tronstad, L.M., Flecker, A.S., Ulseth, A.J., 2007. Improving the fluorometric ammonium method: matrix effects, background fluorescence, and standard additions. J. North Am. Benthol. Soc. 26, 167–177. https://doi.org/10.1899/0887-3593(2007)26[167:itfamm]2.0.co;2

Tonello, M.S., Hebner, T.S., Sterner, R.W., Brovold, S., Tiecher, T., Bortoluzzi, E.C., Merten, G.H., 2019. Geochemistry and mineralogy of southwestern Lake Superior sediments with an emphasis on phosphorus lability. J. Soils Sediments 1–14. https://doi.org/10.1007/s11368-019-02420-5

Twiss, M.R., Rattan, K.J., Sherrell, R.M., McKay, R.M.L., 2004. Sensitivity of phytoplankton to copper in Lake Superior, in: Journal of Great Lakes Research. Elsevier, pp. 245–255. https://doi.org/10.1016/S0380-1330(04)70389-5

Wacklin, P., Hoffmann, L., Komárek, J., 2009. Nomenclatural validation of the genetically revised cyanobacterial genus Dolichospermum (Ralfs ex Bornet et Flahault) comb, nova. Fottea 9, 59–64. https://doi.org/10.5507/fot.2009.005

Walz, H., 2003. Phytoplankton Analyzer Phyto-PAM and Phyto-Win Software V 1.45, System Components and Principles of Operation.

Welschmeyer, N.A., 1994. Fluorometric analysis of chlorophyll a in the presence of chlorophyll b and pheopigments. Limnol. Oceanogr. https://doi.org/10.4319/lo.1994.39.8.1985

Wetzel, R.G., Likens, G.E., 2000. Limnological Analyses, Limnological Analyses. Springer New York. https://doi.org/10.1007/978-1-4757-3250-4

Yurista, P., Kelly, J.R., Miller, S.E., 2011. Lake Superior: Nearshore variability and a landscape driver concept. Aquat. Ecosyst. Heal. Manag. 14, 345–355. https://doi.org/10.1080/14634988.2011.624942

