## Supplementary Materials for "Fluvial seeding of cyanobacterial blooms in oligotrophic Lake Superior"

Chlorophyll µg/L

|  | Blue-green | Green | Brown |
| --- | --- | --- | --- |
| High N:P Microcystis | 262 | 92 | 0 |
| Low N:P Microcysitis | 1379.5 | 0 | 0 |
| High N:P Ankistrodesmus | 0 | 736.5 | 0 |
| Low N:P Ankistrodesmus | 0 | 1246 | 0 |
| High N:P Cyclotella | 63.5 | 0 | 399 |
| Low N:P Cyclotella | 92.5 | 0 | 420 |

Figure S1. PHYTO-PAM measurements for pure cultures used to ground truth measurements. Bars show measurements for each algal group grown in high and low N:P media used during laboratory experiments. Results show that the PHYTO-PAM correctly measures different algal groups.

Table S1. Results of one-way ANOVA with Tukey HSD and Kruskal Wallis tests for differences in site parameters among waterbody types in 2018. No F-value provided for Kruskal-Wallis tests (PP, NH_3_, N:P, and Chl-a), and no p-values provided for Kruskal-Wallis multiple comparisons.

| Parameter | F value | p-value | Comparison | p-value | Significant (p<0.05) |
| --- | --- | --- | --- | --- | --- |
| TOC | 5.7 | 0.0094 | Lake/Pond-Coastal | 0.96 |  |
|  |  |  | River-Coastal | 0.31 |  |
|  |  |  | River-Lake/Pond | 0.0080 | * |
| TN | 0.48 | 0.50 | - | - |  |
| TP | 1.2 | 0.31 | - | - |  |
| POC | 2.6 | 0.093 | - | - |  |
| PON | 2.6 | 0.093 | - | - |  |
| PP | - | 0.75 | - | - |  |
| DOC | 5.7 | 0.0095 | Lake/Pond-Coastal | 0.95 |  |
|  |  |  | River-Coastal | 0.31 |  |
|  |  |  | River-Lake/Pond | 0.0081 | * |
| TDN | 6.2 | 0.0067 | Lake/Pond-Coastal | 1.0 |  |
|  |  |  | River-Coastal | 0.19 |  |
|  |  |  | River-Lake/Pond | 0.0067 | * |
| TDP | 1.3 | 0.30 | - | - |  |
| NH_3_ | - | 0.70 | - | - |  |
| NO_3_^-^ | 3.2 | 0.090 | - | - |  |
| SRP | 0.15 | 0.86 | - | - |  |
| N:P | - | 0.053 |  |  |  |
| Temp | 5.6 | 0.011 | Lake/Pond-Coastal | 0.95 |  |
|  |  |  | River-Coastal | 0.32 |  |
|  |  |  | River-Lake/Pond | 0.0093 | * |
| Chl-a | - | 0.00095 | Lake/Pond-Coastal | - |  |
|  |  |  | River-Coastal | - | * |
|  |  |  | River-Lake/Pond | - | * |
| EC25 | 3.4 | 0.052 | - | - |  |
| Cyanobacterial growth rate (d^-1^) | 1.9 | 0.17 | - | - |  |

Table S2. Results of the paired two-sided t-test for differences in site parameters and laboratory cyanobacterial growth rates between July and August.

| Parameter | t-value | p-value | Significant (p<0.05) |
| --- | --- | --- | --- |
| DOC (µM) | 8.6314 | 5.7 x 10^-09^ | * |
| TDN (µM) | 3.27 | 3.1 x 10^-3^ | * |
| NH_3_ (µM) | 2.11 | 0.045 | * |
| NO_3_ (µM) | 6.0218 | 1.1 x 10^-5^ | * |
| TDP (µM) | 3.38 | 2.4 x 10^-3^ | * |
| SRP (µM) | 2.53 | 0.018 | * |
| TOC (µM) | 7.96 | 2.8 x 10^-8^ | * |
| TN (µM) | 2.3 | 0.030 | * |
| TP (µM) | 2.29 | 0.031 | * |
| N:P (µM) | -0.9 | 0.36 |  |
| chl-a (µg/L) | 1.195 | 0.24 |  |
| POC (µM) | -1.19 | 0.067 |  |
| PON (µM) | -1.19 | 0.067 |  |
| PP (µM) | 0.7 | 0.49 |  |
| Temperature (˚C) | 5.594 | 9.3 x 10^-6^ | * |
| EC_25_ (mS/cm) | -3.474 | 1.9 x 10^-3^ | * |
| Cyanobacterial growth rate (d^-1^) | 0.55 | 0.59 |  |

Table S3. Tabulated results of Kruskal Wallis multiple comparisons.

| Comparison | | Difference |
| --- | --- | --- |
| Group 1 | Group 2 |  |
| 20C-High-Harbor | 20C-High-Lake |  |
| 20C-High-Harbor | 20C-Low-Harbor | * |
| 20C-High-Harbor | 20C-Low-River | * |
| 20C-High-Harbor | 25C-High-Harbor | * |
| 20C-High-Harbor | 25C-High-River |  |
| 20C-High-Harbor | 25C-Low-Harbor | * |
| 20C-High-Harbor | 25C-Low-Lake |  |
| 20C-High-Harbor | 25C-Low-River | * |
| 20C-High-Lake | 20C-Low-Harbor | * |
| 20C-High-Lake | 20C-Low-River | * |
| 20C-High-Lake | 25C-High-Harbor |  |
| 20C-High-Lake | 25C-High-River |  |
| 20C-High-Lake | 25C-Low-Harbor | * |
| 20C-High-Lake | 25C-Low-Lake |  |
| 20C-High-Lake | 25C-Low-River | * |
| 20C-Low-Harbor | 20C-Low-River |  |
| 20C-Low-Harbor | 25C-High-Harbor | * |
| 20C-Low-Harbor | 25C-High-River | * |
| 20C-Low-Harbor | 25C-Low-Harbor | * |
| 20C-Low-Harbor | 25C-Low-Lake | * |
| 20C-Low-Harbor | 25C-Low-River | * |
| 20C-Low-River | 25C-High-Harbor | * |
| 20C-Low-River | 25C-High-River | * |
| 20C-Low-River | 25C-Low-Harbor | * |
| 20C-Low-River | 25C-Low-Lake | * |
| 20C-Low-River | 25C-Low-River | * |
| 25C-High-Harbor | 25C-High-River | * |
| 25C-High-Harbor | 25C-Low-Harbor | * |
| 25C-High-Harbor | 25C-Low-Lake | * |
| 25C-High-Harbor | 25C-Low-River | * |
| 25C-High-River | 25C-Low-Harbor | * |
| 25C-High-River | 25C-Low-Lake |  |
| 25C-High-River | 25C-Low-River | * |
| 25C-Low-Harbor | 25C-Low-Lake | * |
| 25C-Low-Harbor | 25C-Low-River | * |
| 25C-Low-Lake | 25C-Low-River | * |
